# The most primitive extant ancestor of organisms and discovery of definitive evolutionary equations

**DOI:** 10.1101/371575

**Authors:** Kenji Sorimachi

## Abstract

Organisms are classified into three domains, Prokaryota, Archaea, and Eukaryota, and their evolutionary divergence has been characterized based on morphological and molecular features using rationale based on Darwin’s theory of natural selection. However, universal rules that govern genome evolution have not been identified. Here, a simple, innovative approach has been developed to evaluate biological evolution initiating the origin of life: whole genomes were divided into several fragments, and then differences in normalized nucleotide content between nucleotide pairs were compared. Based on nucleotide content structures, *Monosiga brevicollis* mitochondria may be the most primitive extant ancestor of the species examined here. The two normalized nucleotide contents are universally expressed by a linear regression line, (X − Y)/(X + Y) = a (X − Y) + b, where X and Y are nucleotide contents and (a) and (b) are constants. The value of (G + C), (G + A), (G + T), (C + A), (C + T) and (A + T) was ~0.5. Plotting (X − Y)/(X + Y) against X/Y showed a logarithmic function (X − Y)/(X + Y) = a ln X/Y + b, where (a) and (b) are constant. Nucleotide content changes are expressed by a definitive equation, (X − Y) ≈ 0.25 ln(X/Y).

## Introduction

Primitive organisms might appear after long periods of chemical evolution, during which various organic compounds were accumulated. Recently, it was reported that vesicles consisting of lipids and polynucleotides spontaneously replicated under experimental conditions [1]. These pseudo-organisms were the first living things to appear on Earth; however, we cannot trace their origins because the original assembly of chemical compounds was not stable. Only those primitive organisms that established a method of self-replication could survive and continue to evolve. Therefore, it is difficult to identify the true primitive ancestors of life on Earth, as their very make-up would probably be unstable in the current environmental conditions. Although fossilized traces of very early organisms have been found in sedimentary rocks dating from 3.1–3.7 billion years ago [2-5], they do not appear to be the origin of life. It is likely that only primitive organisms that had cell walls could leave a fossil record, which may rule out finding any traces of simpler organisms.

Several key eukaryotic organelles originated from symbioses between separate single-celled organisms [6,7]. For example, mitochondria developed from the proteobacterium *Rickettsia* or its relatives [8,9]. The *Reclinomonas americana* (Protist) mitochondrion (~70 kb), consisting of 97 genes, is thought to be an ancestral mitochondrial DNA (mtDNA), while vertebrate mtDNA (~16 kb), consisting of 37 genes, seems to be constructed with only essential genes for respiration reactions. Based on 13 respiratory genes, the amino acid composition and gene patterns within complete mitochondrial genomes are almost identical among all animal species, except Amoebazoa [10]. Irreversible evolutionary divergence accompanied by increasing G and C content means that the G and C content of descendants should be identical to or higher than that of the ancestor. However, it was shown that the G and C values for *R. americana* (G: 0.148, C: 0.114) mtDNA were higher than those of *Monosiga brevicollis* (G: 0.081, C: 0.059), which were the lowest among the samples examined [11]. In addition, nucleotide regression lines for the two species differed from each other [11,12]. Thus, we concluded that mitochondria might derive directly from primitive organisms [13]. However, there is no scientific evidence that mitochondria are the most primitive extant ancestor of all life. Based on Charles Darwin’s theory of natural selection, all organisms have a single origin and common ancestor, therefore, we predict that the amino acid compositions of their complete genomes should naturally reflect their evolution from bacteria to *Homo sapiens* [14,15].

The two nucleotide relationships were expressed by linear regression lines, which crossed at a single point [11,12]. Thus, we concluded that the origin of life was single and pluripotent. Assuming this conclusion, at the crossing point representing the origin of life, (G ≈ C) and (T ≈ A) to satisfy Chargaff’s parity rules [16,17]: (G – C) ≈ 0 and (T – A) ≈ 0. The two regression lines representing high and low C/G animal mitochondria must satisfy (G ≈ C) and (T ≈ A) at the crossing point for the line based on Chargaff’s parity rule. Therefore, differences in (G – C) and (T – A) reflect mitochondrial evolution. In addition, to satisfy Chargaff’s second parity rule [17], the nucleotide content of the first half of the DNA strand should be equal to that of the second half [18], providing a symmetry between the 5□ and 3□ ends. As animal mitochondrial genomes deviate from Chargaff’s second parity rule [17], the divergence in animal mitochondria increases the unsymmetrical nucleotide content. To detect intragenomic alterations based on increases in the unsymmetrical nucleotide content, a method that investigates nucleotide content differences, i.e. (G – C), (G – A), (G – T), (C – A), (C – T) and (A – T) in the sequentially divided genomes, was developed in the present study. These exciting results could not have been obtained from sequence analyses.

## Materials and methods

All genome sequences were obtained from the National Center for Biotechnology Information GenBank database (http://www.genome.jp/ja). The nucleotide contents of all genomes, which are listed in Extended Data Figs, were normalized (G + C + A + T = 1) because normalized values are independent of species and genome sizes [19,20]. A whole genome was divided into several fragments to evaluate intra-genome alterations due to evolution, and the differences between the two nucleotide contents of six pairs ((G – C), (G – T), (G – A), (C – T), (C – A), (T – A)) were calculated for all fragments. In addition, mitochondria were divided into three equal-size fragments. To easily understand evolution at a glance, these six nucleotide differences were graphically expressed with a single pattern. This appears to be a novel approach. Calculations were carried out with a personal computer, TOSHIBA dynabook T552 (Tokyo, Japan), Windows 10 installed.

## Results and discusion

### Primitive ancestor

Chargaff’s parity rules, where (G = C) and (T = A), were the original rules for inter- [16] and intra- [17] molecular relationships of DNA, and are applicable to chromosomal DNA [21-23]. The normalized nucleotide contents (C, T and A), predicted from complete Genomes, are expressed by the content of the fourth nucleotide (G) using linear regression lines in chromosomal [21], non-animal mitochondrial and chloroplast DNA [22,23]. Classifying animal mitochondria into high and low C/G groups, the content of the first three nucleotides could be similarly expressed by the forth nucleotide content [23], although animal mitochondria deviated from Chargaff’s second parity rule [24,25] (Fig 1*a*). Vertebrate mitochondria overlapped with the high C/G invertebrates, which have high C content (Fig 1*a*), indicating that both groups were descended from the same origin [12,13], and that more highly evolved organisms seem to have a greater cytosine content [11]. Non-animal mitochondria and chloroplasts obeyed Chargaff’s rules[23]. C contents in cellular organelles and chromosomal DNA vs G contents were expressed with linear regression lines, which crossed at a single point (Fig 1*a*). This is consistent with our previous findings [11,12]. In addition, we found that nucleotide content relationships in viral DNA were heteroskedastic (Fig 1*b*).

**Fig. 1.**
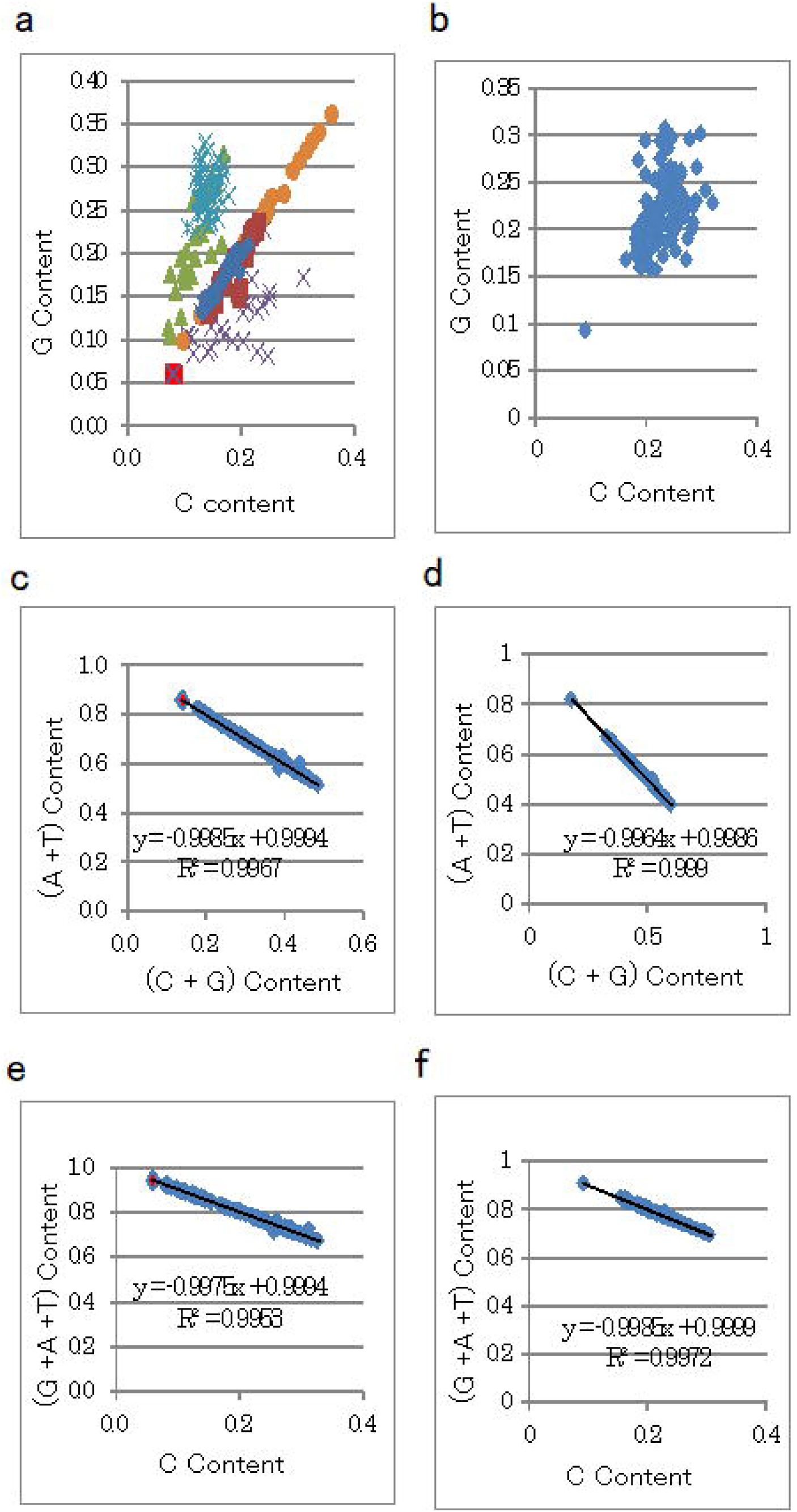
Regression lines. Left and right panels represent cellular organelles and viruses, respectively. Upper panels (a and b): horizontal and vertical axes represent G and C contents, respectively. Upper left panel: C content plotted against G content in vertebrate mitochondria (asterisk), high C/G invertebrate mitochondria (triangle), low C/G invertebrate mitochondria (cross), bacteria (circle), non-animal mitochondria, chloroplasts (diamond), and chromosomes (square). Red square represents *Monosiga brevicollis* mitochondrion. Right upper panel (a): C content plotted against G content in viruses. Middle panels (c and d): Horizontal and vertical axes represent (G + C) and (A + T) contents, respectively. Lower panels (e and f): Horizontal and vertical axes represent (C) and (G + A + T) contents, respectively.

Based on the normalization of the four nucleotides (G + C + T + A = 1), the GC content, (G + C), is expressed by {1 – (A + T)}, where GC content (G + C) and AT content (A + T) are completely linear, not only in chromosomal, non-animal mitochondrial, and chloroplast DNA, but also in animal mitochondrial DNA (Fig. 1c). The same result was obtained for viral DNA (Fig 1d). Assuming irreversible divergence, it is generally thought that GC content increases along with biological evolution. Thus, the organelles that have the lowest GC content might be the most primitive. In the current study, the GC content of the mtDNA of the choanozoan *M. brevicollis* was the lowest among the samples examined. The GC content of the bacterium *Streptomyces coelicolor* was the highest of all the samples, although this organism was not the most evolved. Amongst the viruses examined, the GC content of *Melanoplus sanguinipes* Entomopoxvirus was the lowest, while that of *Papio ursinus* Cytomegalovirus was the highest (Fig 1d).

A previous study [11] stated that the C content of complete mitochondrial genomes reflects biological evolution better than the GC content. Based on the normalization of the four nucleotides, G + C + A + T = 1, C = 1 – (G + A + T). Thus, C content and (G + A + T) are linear. In the current study, the lowest C content was observed in *M. brevicollis* mtDNA, while the highest C content was found in mtDNA from avian species *Gallus gallus* and *Taeniopygia guttata*, and primates *H. sapiens*, *Pan paniscus*, *Pan troglodytes*, and *Gorilla gorilla* (Fig 1e). Amongst the viral genomes, the C content of *Mollivirus sibericum* was the lowest, while that of De-Brazza’s monkey virus was the highest (Fig 1f).

### Genome evolution

The complete genome is represented by four nucleotide contents based on more than a certain amount of randomly chosen fragments, as well as on completely linear fragments [26,27]. Therefore, when the whole genome of bacterium *Ureaplasma urealyticum* (G; 0.131, C; 0.127) was sequentially divided into nine equal fragments, the amounts of the four nucleotides in each fragment were quite similar (Fig 2*a*). This is consistent with our previous results, which indicated that a whole genome may be constructed from small units with similar amino acid compositions [26,27]. Nucleotide content fragments 5 and 6 of the *U. urealyticum* genome (Fig 2*b* and 2*c*). The ratios of (G – C)/(G + C) and (A – T)/(A + T) are called the GC and AT skew, respectively [28]. The skew seems to be based on differences in replication processes between the leading and lagging strands^27^. In particular, replication of the lagging strand increases the probability of mutations as a result of the deamination of cytosine, and the inversion of nucleotide content differences reflects biological divergence. Similar phenomena are observed in mitochondria, which consist of heavy (H) and light (L) chains [30-32]. Plotting the GC skew vs. G content was used to classify animal mitochondria into two groups: high and low C/G^11^. In *M. brevicollis* mitochondria, the nine DNA fragments showed almost the same nucleotide contents (Fig. 2*d*), as was observed in *U. urealyticum* (Fig 2*a*). However, GC and AT content difference inversions were not observed in *M. brevicollis* mtDNA (G: 0.081, C: 0.059) (Fig 2*e* and 2*f*). Thus, the *M. brevicollis* mitochondrion might be more primitive than the *U. urealyticum* chromosome. These results clearly indicate that nucleotide content differences such as (G – C) and (A – T) reflect biological evolution. Therefore, the other nucleotide content differences, (G – A, G – T, C – A and C – T), were examined to determine whether or not these values reflect biological evolution.

**Fig. 2.**
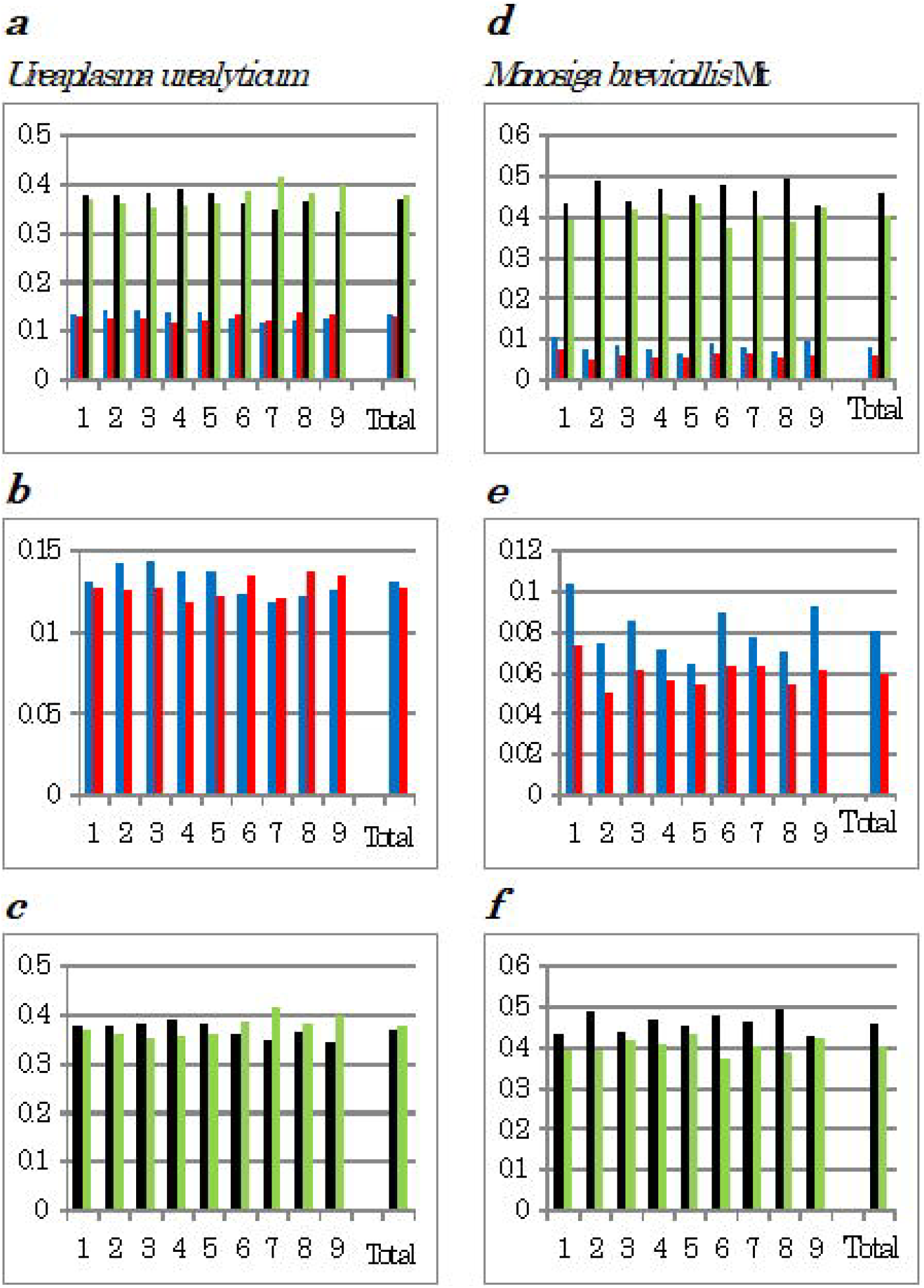
Nucleotide content was sequentially divided into nine fragments. Left side: *Ureaplasma urealyticum* mitochondrion, right side: *Monosiga brevicollis* mitochondrion. G: blue, C: red, A: black, and T: green. Vertical axis represents nucleotide content.

### Organelle evolution

Complete mitochondrial genomes were investigated (Fig 3, left panel). To allow simple visual comparison of inter- and intra-species genome structures, genomes were sequential divided into three fragments throughout subsequent analyses, from which three separate patterns emerged. Six nucleotide content differences were observed among the mitochondria of the four species (*M. brevicollis, P. pallidum, D. discoidium* and *R. americana*) (Fig 3, right panels). The six nucleotide content patterns were conserved within the three fragments among the four species. No inversion of nucleotide content differences was observed in the mtDNA of *M. brevicollis* (G: 0.081, C: 0.059), the mycetozoan *Polysphondylium pallidum* (G: 0.143, C: 0.085), or *Dictyostelium discoideum* (G: 0.171, C: 0.104) (Fig 3), although differences in (G – C) and (T – A) values for *M. brevicollis* mtDNA were the lowest amongst these species. Based on genome sequencing, choanoflagellates are most closely related to animals [33].

As the nucleotide content difference patterns of the three fragments were almost identical for these three species, their nucleotide distributions were judged to be homogeneous, indicating nucleotide content symmetry. Thus, these mitochondria are likely to be primitive. Consistent results were obtained from Ward’s clustering analysis using amino acid compositions predicted from complete mitochondrial genomes as traits^11^. These findings indicate that the *M. brevicollis* mitochondrion is the most primitive among the three. In contrast, AT inversion was observed in the third fragment of *Reclinomonas americana* mtDNA (G: 0.148, C: 0.114), which has previously been proposed as a mitochondrial ancestor [8]. However, differences in (G – C) and (T – A) values in *R. americana* mtDNA were smaller than those in the mtDNA of the previous three organisms. Nucleotide content inversion causes significant differences in nucleotide content patterns as a result of unsymmetrical nucleotide content. Thus, the *R. americana* mitochondrion is probably more evolved than the former three mitochondria. In addition, AT inversion occurred in the following more highly evolved organisms: Mollusca species, squid (*Todarodes pacificus*), octopus (*Octopus vulgaris*), Echinodermata species, sea urchin (*Paracentrotus lividus*), water flea (*Daphnia pulex*), hermit crab (*Pagurus longicarpus*), and Humboldt squid (*Dosidicus gigas*) (S1 Fig). In addition, large positive (G – A) values in the three fragments were observed in *Paragonimus westermani*, while large positive (G – C) and (A – T) values in the three fragments were observed for the mtDNA of representatives of the following phyla: Cnidaria (*Pavona clavus*), Platyhelminthes (*Schistosoma mansoni*), Porifera (*Geodia neptuni*), Arthropoda (*Tigriopus californicus*), and Chordata (*Branchiostoma belcheri*) (S1 Fig.).

**Fig. 3.**
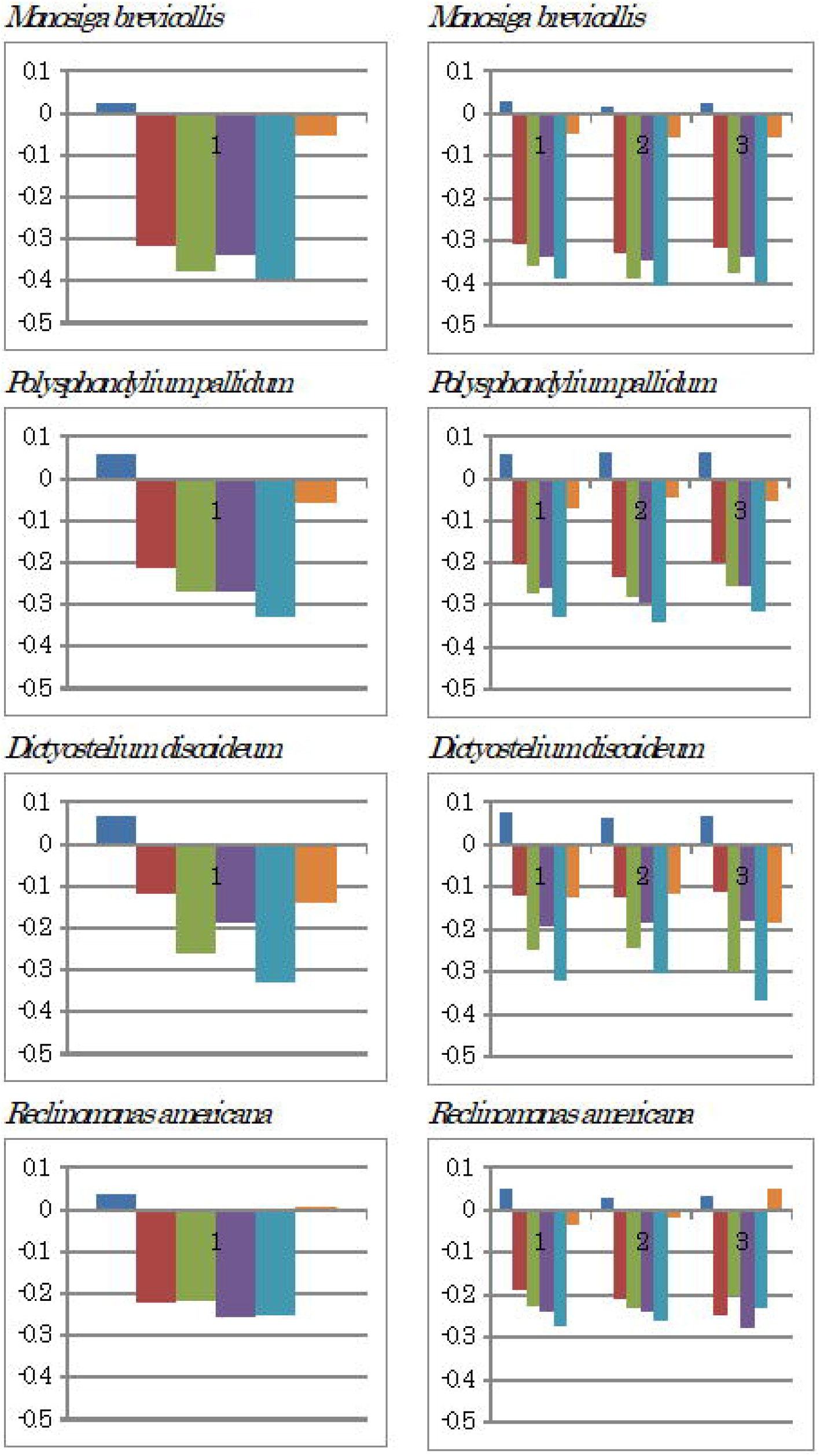
Nucleotide content differences in complete mitochondrial genomes (left side) and the three sequential fragments of each mitochondrial genome (right side). Left to right: (G – C), (G – T), (G – A), (C – T), (C – A) and (T – A).

A positive (C – T) value was characteristically observed in the three fragments of Echinodermata species *Acanthaster planci*, the second and third fragments of Mollusca species *Haliotis rubra*, and in the first fragment of Mollusca species *Lampsilis ornata*. CT inversion occurred in *H. rubra* and *L. ornata* mtDNA (S1 Fig.). The nucleotide content difference patterns of the mtDNA of hemichordates *Saccoglossus kowalevskii* and *Balanoglossus carnosus* differed from each other. Both AT and CT inversions occurred in the first mtDNA fragment of *S. kowalevskii*, while large positive (C – T) and (C – A) value differences occurred in the second and third fragments. AC and AT inversions were observed in *B. carnosus* mtDNA (S1 Fig.). Neither nucleotide inversion nor positive nucleotide differences were observed in the mtDNA of deuterostom, *Xenoturbella bocki*.

In the mtDNA of primate species *H. sapiens*, *P. troglodytes*, *G. gorilla*, *Macaca mulatta*, *Daubentonia madagascariensis*, *Nycticebus coucang*, and *Tupaia belangeri*, nucleotide content difference patterns were quite similar in the first four species, and large positive increases in (C – T) differences in the three fragments clearly indicated evolutionary divergence (Fig. 4). The positive (C – T) differences in all three fragments were characteristic of these four primate mitochondria, while positive increases in (C – T) values were only observed in the third fragment of *N. coucang* and *T. belangeri* mtDNA. In contrast, nucleotide content difference patterns of the prosimian *Lemur catta* completely differed from those of the primates, although TA inversion was observed in the second fragment. The primate mtDNA nucleotide content patterns were also completely different from that of hemichordate *B. carnosus*, although their C contents were the highest among all organisms examined [11]. This finding indicates that mitochondrial structures respect epigenomic evolutionary functions.

**Fig. 4.**
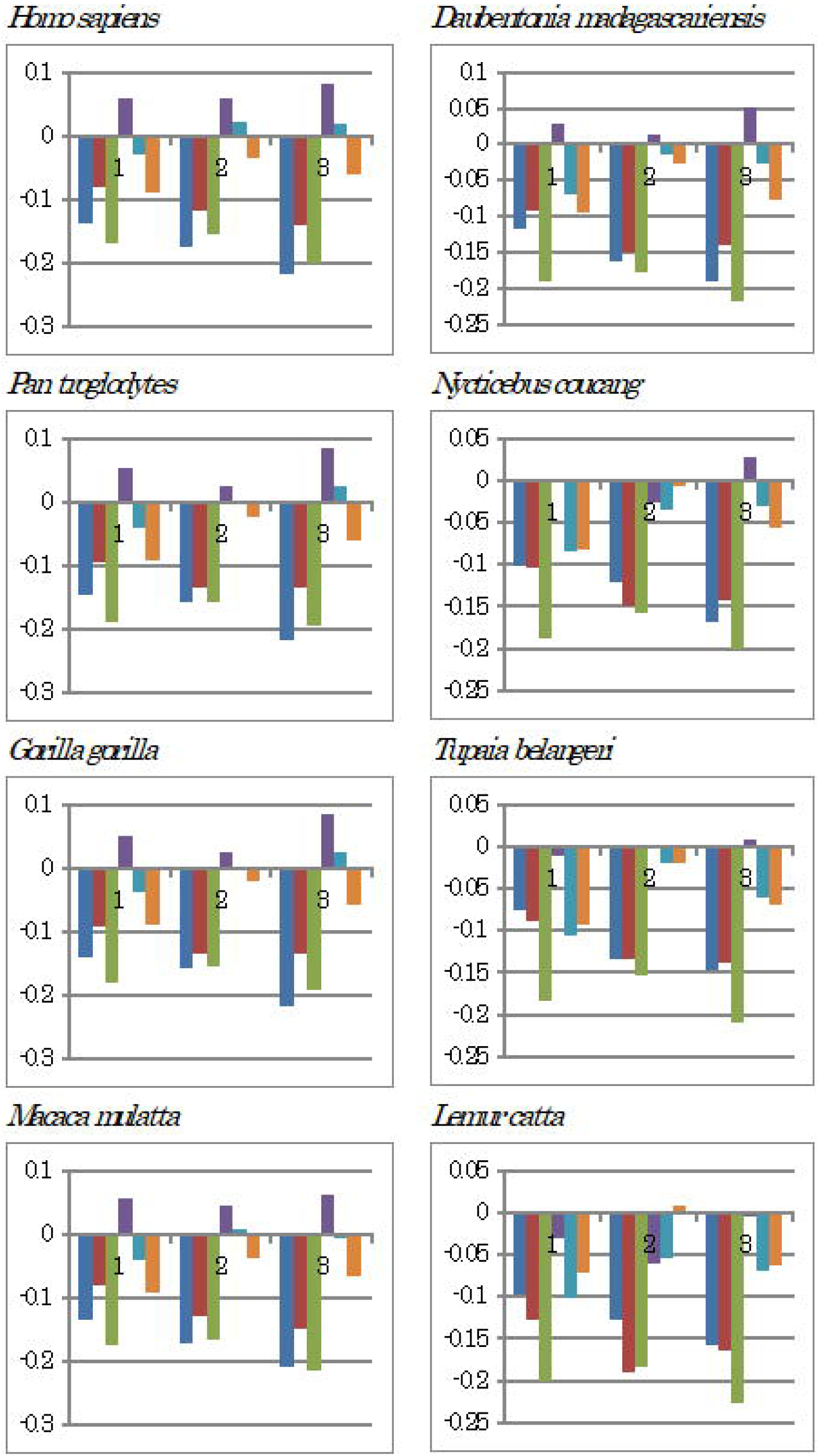
Nucleotide differences in the three fragments of each mitochondrial genome.

Left to right: (G – C), (G – T), (G – A), (C – T), (C – A) and (T – A).

The mitochondria of other vertebrates (S1 Text.), rodents (S2 Fig), ocean-dwelling mammals, cetaceans, aves (S2 Fig.), amphibians (S2 Fig.), reptiles (S2 Fig.), and fishes (S2 Fig.) were also examined. In these organelles, differences in (G – C) and (A – T) or in other nucleotide inversions, as well as GC and AT inversions, increased along with evolution. Consistent results were also obtained from non-animal mitochondria and chloroplasts (S3 Fig.), as well as prokaryotes (S4 Fig.). In the microsporidian protozoan *Encephalitozoon cuniculi*, GC and AT inversions, as well as other nucleotide content inversions, were observed (S5 Fig.).

### Virus evolution

The *M. sanguinipes* Entomopoxvirus genome had the lowest G content among the viruses examined (Fig. 1), and AT inversion was also observed (S6 Fig.). In *P. ursinus* Cytomegalovirus, whose GC content was the highest, both GC and AT inversions occurred. *Mollivirus sibercum*, which had the lowest C content, showed both GC and AT inversions, while DeBrazza’s monkey virus 1 showed both GA and GT inversions.

Ebola haemorrhagic fever, caused by the Ebola virus, can be fatal in humans. The Reston, Sudan, and Zaire strains did not show any nucleotide inversion in the three genome fragments (Fig. 5). In the Tai Forest and Bundibugyo strains, however, CT inversion was clearly observed in the first fragment, and was accompanied by a decrease in GT content difference. These nucleotide content differences corresponded to the GC contents of the strains: Reston (G: 0.198, C: 0.210)[34], Sudan (G: 0.198, C: 0.216)[35], Zaire (G: 0.198, C: 0.213)[36], Tai Forest (G: 0.192, C: 0.231)[37], and Bundibugyo (G: 0.192, C: 0.228)[37]. An increase in C content and decrease in G content were observed in the Tai Forest and Bundibugyo strains. The calculated GC contents for the strains were: 0.406 (Reston), 0.411 (Zaire), 0.414 (Sudan), 0.420 (Bundibugyo), and 0.423 (Tai Forest). These results may indicate that Ebola virus evolution occurred over a short period of time.

**Fig. 5.**
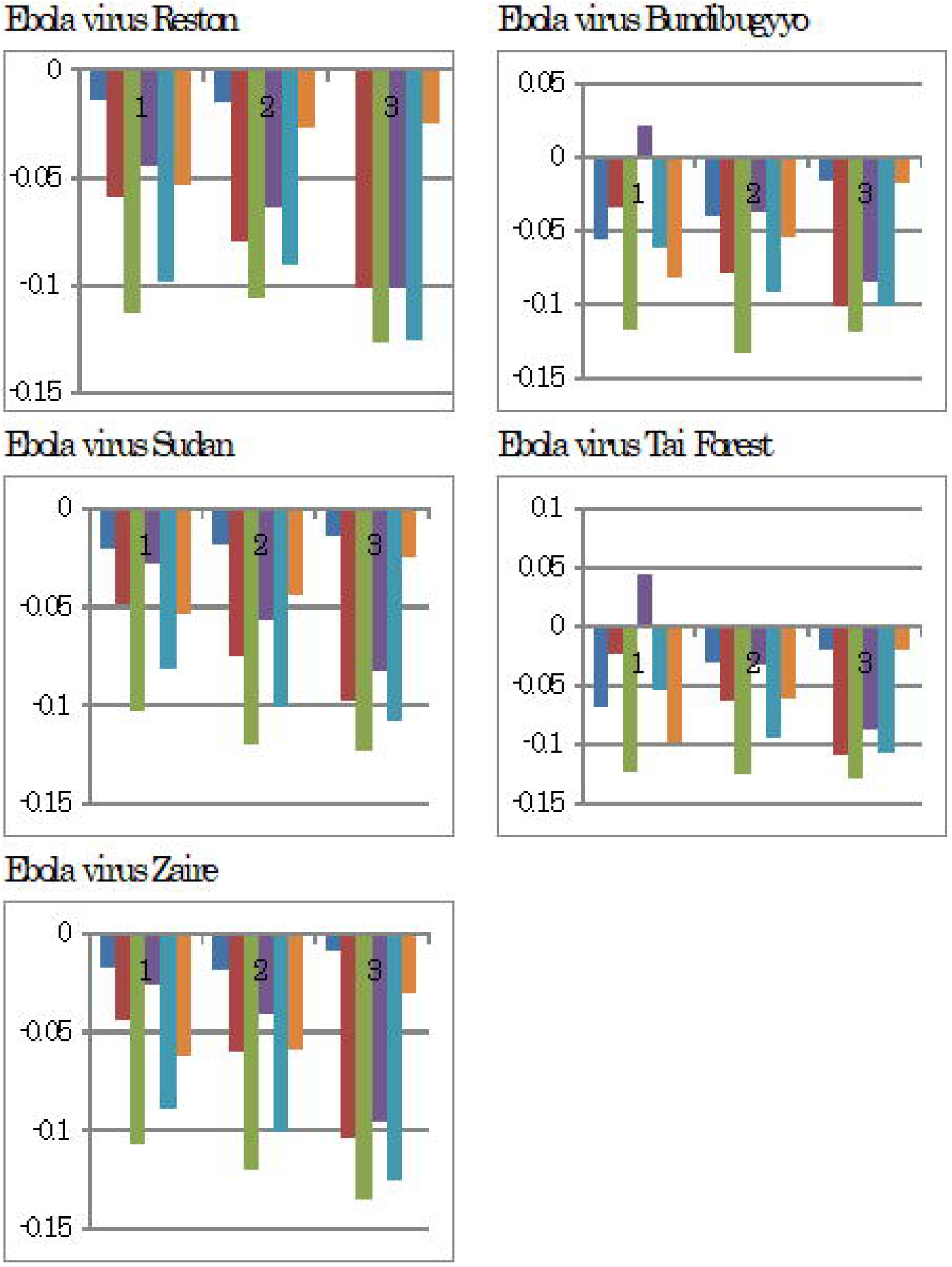
Nucleotide content differences in the three fragments of Ebola virus genomes. Left to right: (G – C), (G – T), (G – A), (C – T), (C – A) and (T – A).

### Interspecies evolution

Nucleotide content differences were calculated in vertebrate mitochondria (S7 Fig.), invertebrate mitochondria (S7 Fig.), non-animal mitochondria (S7 Fig.), chloroplasts (S7 Fig.) and nuclear DNA (S7 Fig.).

### Definitive universal equation

Plotting (X – Y)/(X + Y) against (X – Y), the following linear relationship was obtained in mitochondria, chloroplasts, and chromosomes (Fig.6*a*), and viruses (Fig. 6*b*): (X – Y)/(X + Y) = a (X – Y) + b, where X and Y are nucleotide contents, and (a) and (b) are constants. As (b) was almost null and (a) was ~2.0, (X – Y)/(X + Y) ≈ 2.0 (X – Y). In these genome analyses, which are independent of Chargaff’s parity rules (S8 Fig., left panels), the values of (a) for (G, C), (G, A), (G, T), (C, T), (C, A) and (A, T) were 2.5858, 1.85558, 1.9908, 1.9771, 1.9968 and 1.5689, respectively. Based on these results, (G + C), (G + A), (G + T), (C + A), (C + T) and (A + T) were 0.39, 0.54, 0.50, 0.51, 0.50 and 0.64, respectively. In virus genome analyses (S8 Fig., right panels), the constant values for (a) were 1.9–2.1, and the values for (X + Y) were 0.47–0.53. In contrast, in the normalization of nucleotide contents (G + C + A + T = 1), as (G = C) and (A = T) based on Chargaff’s parity rules, (2G + 2A = 1) is obtained. This equation is altered to (G + A = 0.5). This value is consistent with the value obtained above from genome analyses. Similarly, (G + T = 0.5), (C + A = 0.5) and (C + T = 0.5), although (G + C) and (A + T) cannot be determined. Therefore, the four nucleotide contents are expressed by the following regression lines, plotted against G content: A = 0.5 – G, T = 0.5 – G, C = G and (G = G). Lines G and C overlap, as do lines A and T, and the former line is symmetrical to the latter against line (y = 0.25). The intercepts of lines G and C are close to the origin, while those of lines A and T are close to 0.5 at the vertical and horizontal axes. All organisms from bacteria to *H. sapiens* are located on the diagonal lines of a 0.5 square, termed the “Diagonal Genome Universe”, using the normalized values that obey Chargaff’s parity rules [27]. These relationships lead to (G or C) + (A or T) = 0.5. The present results indicate that a linear regression line equation, (X – Y)/(X + Y) = a (X – Y) + b, universally represents all normalised values, including the values deviating from Chargaff’s parity rules. This newly discovered equation clearly reflects not only Chargaff’s parity rules, based on hydrogen bonding between two nucleotides, but also natural rule.

A linear regression line was not obtained when using randomly chosen value (Fig. 6*c*). Furthermore, plotting (X – Y)/(X + Y) against (X/Y), the following logarithmic function was obtained for all tested genomes as well as when using randomly chosen values (Fig. 6 *a’-c’*): (X – Y)/(X + Y) = a ln (X/Y) + b. As (b) was almost null and (a) was ~0.5, (X – Y)/(X + Y) ≈ 0.5 ln (X/Y). The ratio between two values, (X/Y), can be expressed by a logarithmic function, ~0.5 ln (X/Y) ≈ (X – Y)/(X + Y). Plotting the GC skew vs. G content, animal mitochondria were classified into two groups: high and low C/G [11]. This fact indicates that the ratio C/G and the GC skew are evolutionarily related to each other. Any change can be expressed universally by a definitive logarithmic function, (X – Y)/(X + Y) = a ln (X/Y) + b. The present results indicate that cellular organelle evolution is strictly controlled under these characteristic rules, although non-animal mitochondria, chloroplasts, and chromosomes are controlled under Chargaff’s parity rule [17].

**Fig. 6.**
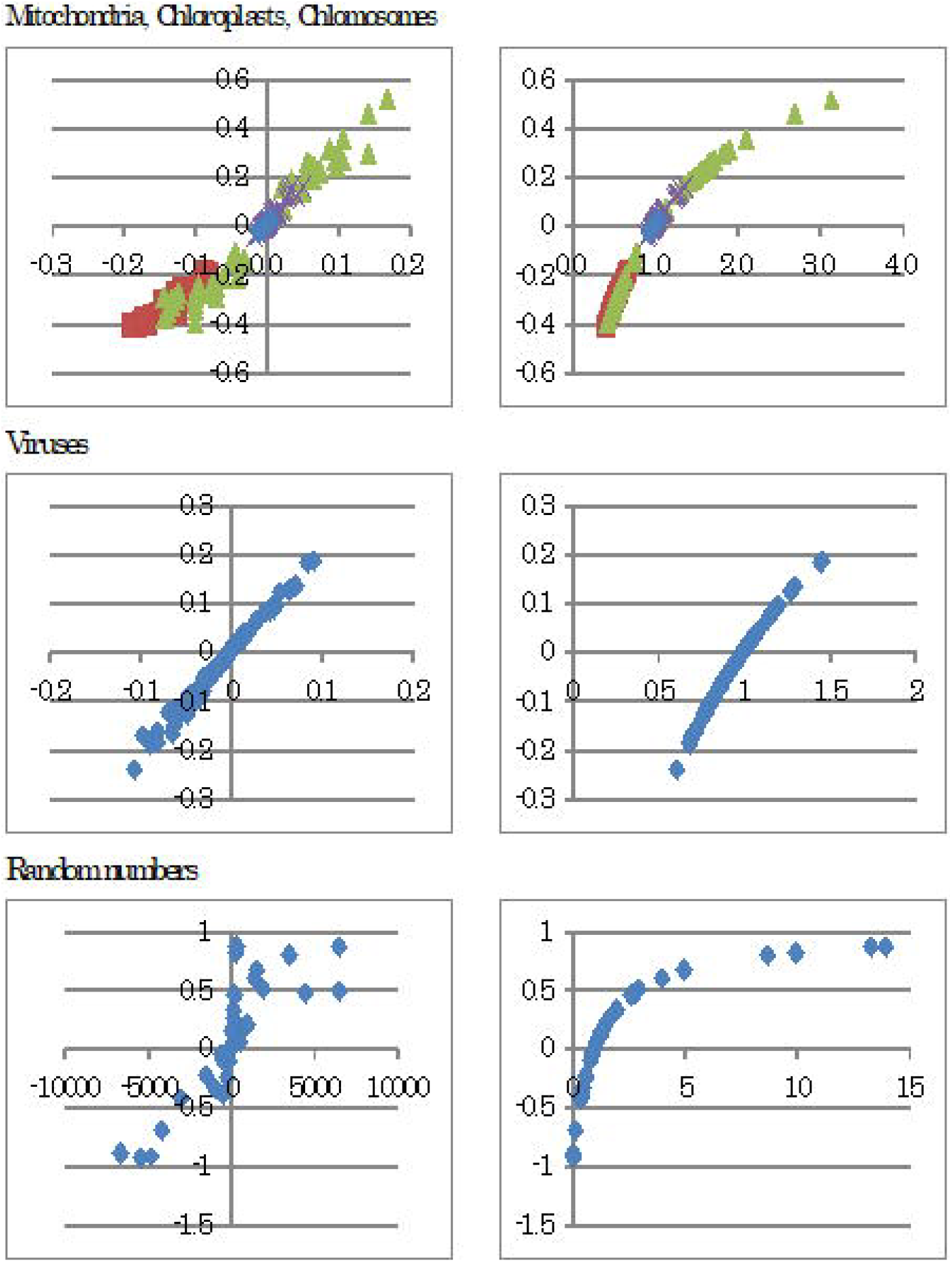
Universal rules. Left side: relationship between (G – C) and (G – C)/(G + C), expressed by a linear regression, and right side: relationship between (G/C) and (G – C)/(G + C), expressed by a logarithmic function. The random numbers were not normalized.

## Conclusions

Our findings showed that the most primitive extant ancestor of all living organisms is the *M. brevicollis* mitochondrion. In the normalised genome values, the relationship between the nucleotide difference, (X – Y), and (X – Y)/(X +Y), including biological evolution, can be expressed definitively by a linear regression line: (X – Y)/(X +Y) = a (X – Y) + b, where (a ≈ 2.0) and (b ≈ 0) are constants. (X + Y) = approximately 0.5. The nucleotide difference, (X – Y), is generally expressed by a linear regression line that crosses “**0**” at (X = Y), representing no evolution. In addition, the relationship between the ratio (X/Y) and (X – Y)/(X +Y) can be expressed definitively by a logarithmic function, (X – Y)/(X + Y) = a ln (X/Y) + b, where (a ≈ 0.5) and (b ≈ 0) are constants. The relationship between the skew “(X – Y)/(X + Y)” and the ratio (Y/X) is universally expressed by a logarithmic line that crosses “1” at (X = Y), representing no change. As the left sides, (X – Y)/(X + Y), are equal in both equations, **2.0** (X – Y) ≈ **0.5** ln(X/Y). Finally, (X – Y) ≈ **0.25** ln(X/Y) using the normalized values from viruses to *H. sapiens* (Fig. 7).

**Fig. 7.**
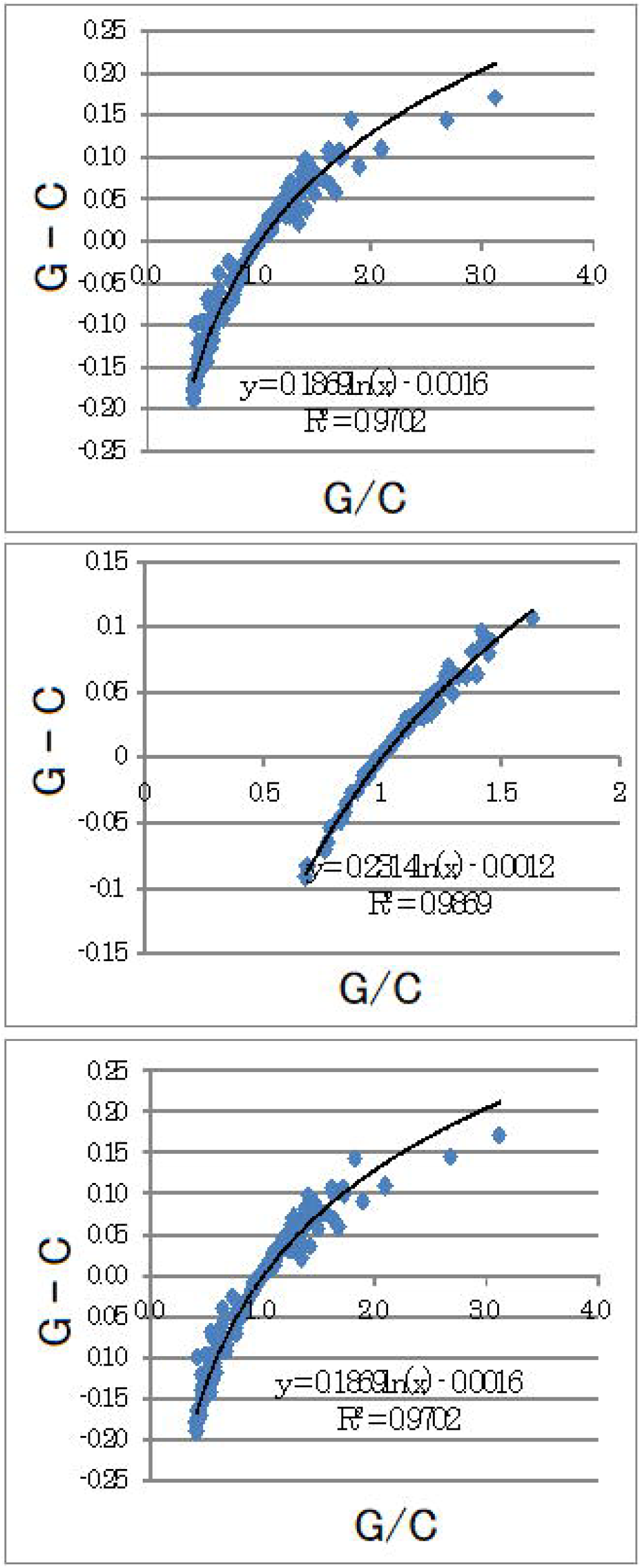
Universal rule governing genome evolution from viruses to *H. sapiens*. Upper pane: cellular organelles, middle panel: viruses, and lower panel: cellular organelles plus viruses.

## Supporting information

**S1 Fig. Nucleotide content differences in the three fragments of the mitochondrial genomes of invertebrates.** Left to right: (G – C), (G – T), (G – A), (C – T), (C – A), and (T – A).

**S2 Fig. Nucleotide content differences in the three fragments of the mitochondrial genomes of vertebrates.** Left to right: (G – C), (G – T), (G – A), (C – T), (C – A), and (T – A).

**S3 Fig. Nucleotide content differences in the three fragments of the mitochondrial genomes of non-animal mitochondria and chloroplasts.** Left to right: (G – C), (G – T), (G – A), (C – T), (C – A), and (T – A).

**S4 Fig. Nucleotide content differences in the three fragments of bacterial genomes.** Left to right: (G – C), (G – T), (G – A), (C – T), (C – A), and (T – A).

**S5 Fig. Nucleotide content differences in the three fragments of the chromosome of microsporidian protozoan species *Encephalitozoon cuniculi*.** Left to right: (G – C), (G – T), (G – A), (C – T), (C – A), and (T – A).

**S6 Fig. Nucleotide content differences in the three fragments of viral genomes.** Left to right: (G – C), (G – T), (G – A), (C – T), (C – A), and (T – A).

**S7 Fig. Nucleotide differences.** (a) Mitochondria of aquatic vertebrates (blue arrow) and terrestrial vertebrates (red arrow). (b) Mitochondria of high C/G invertebrates (black arrow) and low C/G invertebrates (red arrow). (c) Non-animal mitochondria of fungi (blue arrow) and plants (green arrow). (d) Chloroplasts. (e) Chromosomes of prokaryotes (blue arrow), archaea (green arrow), and eukaryotes (red arrow).

**S8 Fig. Universal rules.** Left side: relationship between (X – Y) and (X – Y)/(X + Y), and right side: relationship between (X/Y) and (X – Y)/(X + Y). (a): organelles and (b): viruses.

**S1 Table. List of viruses used.**

**S1 Text. Application of newly developed analytical method to various organisms’ mitochondrial genomes.**

